# Synergistic interaction of caspofungin combined with posaconazole against *FKS* wild-type and mutant *Candida auris* planktonic cells and biofilms

**DOI:** 10.1101/2022.08.31.506097

**Authors:** Noémi Balla, Fruzsina Kovács, Bence Balázs, Andrew M Borman, Aliz Bozó, Ágnes Jakab, Zoltán Tóth, László Majoros, Renátó Kovács

## Abstract

The *in vitro* efficacy of caspofungin against *FKS* wild type and mutant *Candida auris* isolates was determined in the presence of posaconazole. Drug–drug interactions were assessed utilizing the fractional inhibitory concentration indices (FICIs), the Bliss independence model and a LIVE/DEAD viability assay. Median planktonic minimum inhibitory concentrations (pMICs) of *C. auris* isolates were between 0.5 and >2 mg/L for caspofungin and between 0.125 and >0.25mg/L for posaconazole. Median pMICs for caspofungin and posaconazole in combination showed a 4- to 256-fold decrease compared to caspofungin and a 2- to 512-fold decrease compared to posaconazole alone. The median sessile minimum inhibitory concentrations (sMICs) of isolates ranged from 32 to >32 mg/L and from 0.06 to >2 mg/L for caspofungin and posaconazole, respectively. Median sMICs for caspofungin and posaconazole in combination showed an 8- to 128-fold decrease compared to caspofungin and a 4- to 512-fold decrease compared to posaconazole alone. Caspofungin and posaconazole showed a synergistic interaction, especially against sessile cells (FICI from 0.033–0.375 and 0.091–0.5, and Bliss cumulative synergy volumes were 6.96 and 32.39 for echinocandin-susceptible and -resistant isolates, respectively). In line with the checkerboard-based findings, synergistic interactions were confirmed by a fluorescent microscopic LIVE/DEAD viability assay. The caspofungin-exposed (4 mg/L) *C. auris* biofilms exhibited increased cell death in the presence of posaconazole (0.03 mg/L) compared to untreated, caspofungin-exposed and posaconazole-treated sessile cells. The disrupted biofilm structure and increase in cell death was observed for both echinocandin-susceptible and echinocandin-resistant isolates. Despite the favourable effect of caspofungin in the presence of posaconazole, further *in vivo* studies are needed to confirm the clinical therapeutic potential of this combination when treating *C. auris*.

**Contribution to the field:** *Candida auris* is an emerging fungal pathogen, presumably related to global warming, which is associated with nosocomial infections and is considered a serious health threat worldwide. The treatment of *C. auris* infections is challenging due to the high level of drug resistance against the traditional antifungal agents. Given the low frequency of resistance to echinocandins, they are recommended as first-line therapy for the management of *C. auris* infections; however, treatment is complicated by the development of resistance in patients receiving long-term echinocandin treatment. In addition, the biofilm forming ability of this species further complicates the echinocandin-based therapeutic strategies. Combination-based approaches using existing drugs are viable alternatives to overcome the difficult-to-treat *C. auris-related* infections, including biofilm associated cases. In this study, we examined the *in vitro* efficacy of caspofungin and posaconazole against *FKS* wild-type and mutant *C. auris* planktonic cells and biofilms using classic checkerboard-based investigations and fluorescent imaging. Based on our results, the efficacy of caspofungin and posaconazole is unquestionable, having been confirmed against biofilms, especially in the case of *FKS* mutants at clinically achievable and safe drug concentrations. This study suggests that the administration of caspofungin with posaconazole may help to expand potential treatment strategies.

## 1. Introduction

*Candida auris* is an emerging pathogen, presumably related to global warming, and causes invasive infections and nosocomial outbreaks worldwide (Casadewall *et al.* 2021). The Centers for Disease Control and Prevention (CDC) have expressed alarm that more than 90% of isolates are resistant to fluconazole, frequently accompanied by a decreased susceptibility to amphotericin B (30% resistance) and echinocandins (3 to 7% resistance) (Kordalewska *et al.* 2018, Lockhart 2019, Zhu *et al.* 2020). Based on current data derived from New York-New Jersey, echinocandin resistance increased from 0% to 4% between 2016 and 2020 (Kilburn *et al*. 2022). Furthermore, a worrisome 37% increase in minimum inhibitory concentrations (MICs) to caspofungin was reported in a multicenter analysis derived from India (Kathuria *et al.* 2015). Moreover, the majority of *C. auris* isolates are capable of biofilm development on a variety of surfaces, promoting nosocomial transmission and a higher ratio of clinical failures (Kean *et al*. 2018, Horton *et al*. 2020). It is noteworthy that indwelling devices were the source of nearly 90% of *C. auris* bloodstream infections, emphasizing the clinical importance of these sessile communities (Sayeed *et al*. 2020, Horton *et al*. 2020). Although echinocandins have good activity against biofilms, their efficacy is significantly lower against *C. auris* than against *Candida albicans* planktonic cells or biofilms (Sherry *et al*. 2017). Nevertheless, given the relatively low frequency of resistance to echinocandins, they are recommended as first-line agents for the treatment of invasive *C. auris* infections; however, treatment is complicated by the development of resistance in patients receiving long-term echinocandin treatment (Cortegiani *et al.* 2018, Aldejohann *et al.* 2022). Several investigators have proposed combination-based therapeutic approaches using existing drugs to overcome the difficult-to-treat *C. auris-*related infections, including biofilm associated cases, increasing the likelihood of therapeutic success (Bidaud *et al*. 2020, Brennan-Krohn *et al.* 2021, Caballero *et al.* 2021, Nagy *et al.* 2021, Vitale 2021). Based on previously published results, the combination of caspofungin and posaconazole has shown high efficacy against both *C. albicans* and *Candida glabrata* echinocandin-susceptible and -resistant isolates (Chen *et al.* 2013, Denardi *et al.* 2017, Khalifa *et al.* 2021). However, whether combinations of posaconazole with echinocandins possess synergistic interactions against *C. auris*, especially against biofilms, has been poorly studied. A study published by O’brien *et al.* (2020) examined only one posaconazole (1 mg/L) and caspofungin (4 mg/L) combination against planktonic *C. auris* cells, and the obtained results were controversial. Hence, the aim of this study was to examine the *in vitro* efficacy exerted by caspofungin and posaconazole combinations against echinocandin-susceptible (wild type) and echinocandin-resistant (*FKS* mutant) *C. auris* planktonic cells and biofilms.

## 2. Materials and methods

### 2.1. Isolates

Eight isolates were used, each belonging to a distinct clade of *C. auris,* referred to as South Asian, East Asian, South African and South American clades. The characteristics of the isolates are presented in Table 1. Four out of eight isolates were *FKS* mutant, with elevated MIC values to caspofungin. The tested isolates were identified to the species level by matrix-assisted laser desorption/ionisation time-of-flight mass spectrometry (MALDI-TOF). Clade delineation was conducted by polymerase chain reaction (PCR) amplification and sequencing of the 28S ribosomal DNA gene and the internal transcribed spacer region 1, as described previously. Biofilm formation was assessed with the crystal violet assay, as previously described by Kovács *et al.* (2016).

**Table 1.** Characteristics of *Candida auris* isolates used in this study.

| Isolates | Clade | Isolation Source | *FKS* mutations |
| --- | --- | --- | --- |
| Ca_1 | South Asian | wound swab | HS1 WT<br>HS2 R1354H |
| Ca_2 | South Asian | perianal swab | HS1 WT<br>HS2 R1354H |
| Ca_3 | South Asian | Central line | HS1 S639Y<br>HS2 WT |
| Ca_4 | South Asian | wound swab | HS1 S639P<br>HS2 WT |
| Ca_5 | South Asian | Unknown | HS1 WT<br>HS2 WT |
| Ca_6 | East Asian | Unknown | HS1 WT<br>HS2 WT |
| Ca_7 | South African | Tracheostomy | HS1 WT<br>HS2 WT |
| Ca_8 | South American | Blood | HS1 WT<br>HS2 WT |

HS1 corresponds to hot spot 1
HS2 corresponds to hot spot 2
WT corresponds to wild type

### 2.2. Whole genome sequencing of isolates

Library preparation was performed using the tagmentation based Illumina DNAFlex Library Prep kit (Illumina, San Diego, CA, USA), according to the manufacturer’s protocol. Paired - end 300 bp sequencing was executed on an Illumina MiSeq instrument. Raw sequencing reads were aligned to the *C. auris* B8441 reference genome using the Burrows-Wheeler Aligner algorithm. Genetic variants (single nucleotide polymorphisms, mutation, indel variants) were determined using the GATK algorithm. Library preparations, sequencing and data analysis were performed at the Genomic Medicine and Bioinformatics Core Facility of the University of Debrecen, Hungary.

### 2.3. Antifungal susceptibility testing for planktonic cells

The planktonic MIC (pMIC) was determined based on the M27-A3 protocol released by the Clinical Laboratory Standards Institute (CLSI 2008). Susceptibility to caspofungin pure powder (Merck, Budapest, Hungary) and posaconazole pure powder (Merck, Budapest, Hungary) was determined in RPMI-1640 (with L-glutamine and without bicarbonate, pH 7.0, and with MOPS; Merck, Budapest, Hungary). The drug concentrations ranged from 0.0009 to 0.25 mg/L for posaconazole and from 0.03 to 2 mg/L for caspofungin. The pMICs were determined as the lowest antifungal concentration that exerts at least 50% growth inhibition compared with the untreated growth control and are presented as the median value of three independent experiments per isolate. *Candida parapsilosis* ATCC 22019 and *Candida krusei* ATCC 6258 were used as quality control strains.

### 2.4. Biofilm development

One-day-old biofilms were prepared as described previously (Kovács *et al.* 2016, Nagy *et al.* 2021). Briefly, following 48 h culturing on Sabouraud dextrose agar (Lab M Ltd., Bury, United Kingdom), *C. auris* cells were harvested by centrifugation (3000 *g* for 5 min), washed three times in sterile physiological saline, and the final density of the inoculums was adjusted in RPMI-1640 broth to 1 × 10^6^ cells/mL. Afterwards, 100 μL aliquots were inoculated onto flat-bottom 96-well sterile microtitre plates (TPP, Trasadingen, Switzerland) and incubated statically in darkness at 37°C for 24 hours.

### 2.5. Assessment of antifungal susceptibility for biofilms

The caspofungin concentrations for the biofilm MIC (sMIC) determination ranged from 0.5 to 32 mg/L, while the examined posaconazole concentrations ranged from 0.007 to 2 mg/L. Biofilms were washed three times with sterile physiological saline. After incubation at 37°C for 24 hours, biofilms were washed with sterile physiological saline, and an XTT-assay was performed, as described previously (Kovács *et al*. 2016, Nagy *et al*. 2020, Nagy *et al*. 2021). The change (%) in metabolic activity was calculated based on absorbance (*A*_492nm_), as:

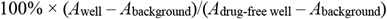

The sMICs were defined as the lowest drug concentration resulting in at least a 50% metabolic activity decrease compared with untreated control cells (Kovács *et al*. 2016, Nagy *et al*. 2020, Nagy *et al*. 2021) and are presented as the median value of three independent experiments per isolate.

### 2.6. Assessment of synergy between caspofungin and posaconazole

Drug-drug interactions between caspofungin and posaconazole were assessed by the two-dimensional checkerboard broth microdilution assay, as previously described (Meletiadis *et al.* 2005, Kovács *et al.* 2016, Nagy *et al.* 2020, Nagy *et al.* 2021, Bidaut *et al.* 2021). Planktonic and sessile cells were prepared with 2 × 10^4^ cells/mL and 1 × 10^6^ cells/mL, respectively, containing different concentrations of each drug combination. The concentrations tested corresponded to the values described in the susceptibility experiments. Afterwards, plates were incubated for 24 hours at 37°C. The data obtained from the checkerboard tests were evaluated by the fractional inhibitory concentration index (FICI), which was expressed as:

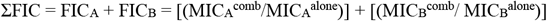

where MIC_A_^alone^ and MIC_B_^alone^ are the MICs of drugs A and B when used alone, and MIC_A_^comb^ and MIC_B_^comb^ are the MICs of drugs A and B in combination at isoeffective combinations, respectively (Meletiadis *et al.* 2005, Kovács *et al.* 2016, Nagy *et al.* 2020, Nagy *et al.* 2021). The FICIs were determined as the lowest ΣFIC. MICs of the drugs alone and those of all isoeffective combinations were determined as the lowest concentration resulting in at least 50% reduction in turbidity and metabolic activity compared with the untreated control cells for planktonic and sessile population, respectively. FICIs were determined in three independent experiments and are expressed as the median value. A synergistic interaction was defined as FICI≤0.5, while 0.5<FICI≤4 was considered to be an indifferent interaction and FICI>4 was considered to be an antagonistic interactions (Meletiadis *et al*. 2005, Kovács *et al*. 2016, Nagy *et al*. 2020, Nagy *et al*. 2021).

To further evaluate the nature of the caspofungin and posaconazole interactions, MacSynergy II analysis was applied, which employs the Bliss independence algorithm in a Microsoft Excel–based interface to assess the nature of interactions (Prichard and Shipman 1990, Rhoden *et al*. 2020, Nagy *et al*. 2020, Nagy *et al*. 2021). This algorithm calculates the difference (ΔE) in the predicted percentage of growth (E_ind_) and the experimentally observed percentage of growth (E_exp_), to define the interaction of the drugs used in combination. The MacSynergy II model expresses interaction volumes and determines positive volumes as synergistic and negative volumes as antagonistic (Prichard and Shipman 1990, Rhoden *et al.* 2020, Nagy *et al*. 2020, Nagy *et al*. 2021). The E values of all combinations are presented on the z-axis in the three-dimensional plot. Synergy or antagonism is significant if the interaction log volumes are >2 or <2, respectively, while log volume values between >2 and 5 correspond to minor synergy, between >5 and 9 shows moderate synergy, >9 shows strong synergy, and the negative values correspond to minor, moderate and strong antagonistic interaction, respectively (Prichard and Shipman 1990, Rhoden *et al.* 2020, Nagy *et al.* 2020, Nagy *et al.* 2021). The synergy volumes were calculated at the 95% confidence level.

### 2.7. Biofilm viability assay

The effect of combinations on viability was examined using the LIVE/DEAD^®^ BacLight™ assay against all isolates tested and pictures form one-one representative echinocandin susceptible (strain Ca_5) and resistant (Ca_1) isolate were presented. One-day-old biofilms were grown on the surface of a 4-well Permanex slide (Lab-Tek^®^ Chamber Slide™ System, VWR, Debrecen, Hungary). The preformed biofilms were washed three times with sterile physiological saline, and various drug concentrations, chosen based on the checkerboard results, were added to the samples as follows: 4 mg/L caspofungin, 0.03 mg/L posaconazole, and 4 mg/L caspofungin combined with 0.03 mg/L posaconazole. Following 24 hours of antifungal treatment, the sessile cells were washed with sterile physiological saline, and the ratio of viable and dead cells was evaluated using the fluorescent LIVE/DEAD^®^ BacLight™ viability kit (ThermoFisher scientific, USA), as described in our previous works (Nagy *et al.* 2020, Kovács *et al.* 2021). Biofilms were exposed with Syto 9 (3.34 mM solution in DMSO) and propidium iodide (20 mM solution in DMSO) for 15 minutes in darkness at 37°C to examine viable and dead *Candida* cells, respectively. Fluorescent cells were studied with a Zeiss AxioSkop 2 mot microscope (Jena, Germany) coupled with a Zeiss AxioCam HRc camera (Jena, Germany). Analysis of the images was performed using Axiovision 4.8.2 (Jena, Germany).

## 3. Results

Whole genome sequencing and *FKS1* analysis was performed for all *C. auris* isolates, and four echinocandin-sensitive isolates presented the wild-types genotype. Four isolates (Ca_1, Ca_2, Ca_3 and Ca_4) were considered to be resistant to caspofungin, based on the tentative MIC breakpoint recommended by the CDC (≥2 mg/L). Two isolates (Ca_1 and Ca_2) contained the R1354H mutation in hot-spot 2 of the *FKS1* gene. Moreover, two well-described S639Y and S639P mutations were observed in the hot-spot 1 region for Ca_3 and Ca_4, respectively (Table 1).

The median and range of MICs for planktonic isolates and *C. auris* biofilms are presented in Table 2. Using the microdilution method, isolates were shown to exhibit pMICs for caspofungin alone from 0.5–1 mg/L and >2 mg/L for echinocandin-susceptible and echinocandin-resistant strains, respectively. In the case of posaconazole, the median pMICs ranged from 0.125 to >0.25 mg/L for both susceptible and resistant strains, respectively. The sMICs for caspofungin alone were from 32 to >32 mg/L, regardless of *FKS* phenotype. The biofilm-forming isolates exhibited sMICs for posaconazole alone from 0.25 to >2 mg/L and from 0.06 to >2 mg/L for echinocandin-susceptible and echinocandin-resistant strains, respectively. In the case of *FKS* mutant isolates, the median pMICs observed in combination showed a 4- to 256-fold reduction for caspofungin and a 2- to 256-fold reduction for posaconazole. The wild type strains showed a reduction in MIC values for posaconazole (2- to 512-fold), while a 0 to 2-fold increase was observed in caspofungin MICs in combination with posaconazole (Table 2).

**Table 2.**
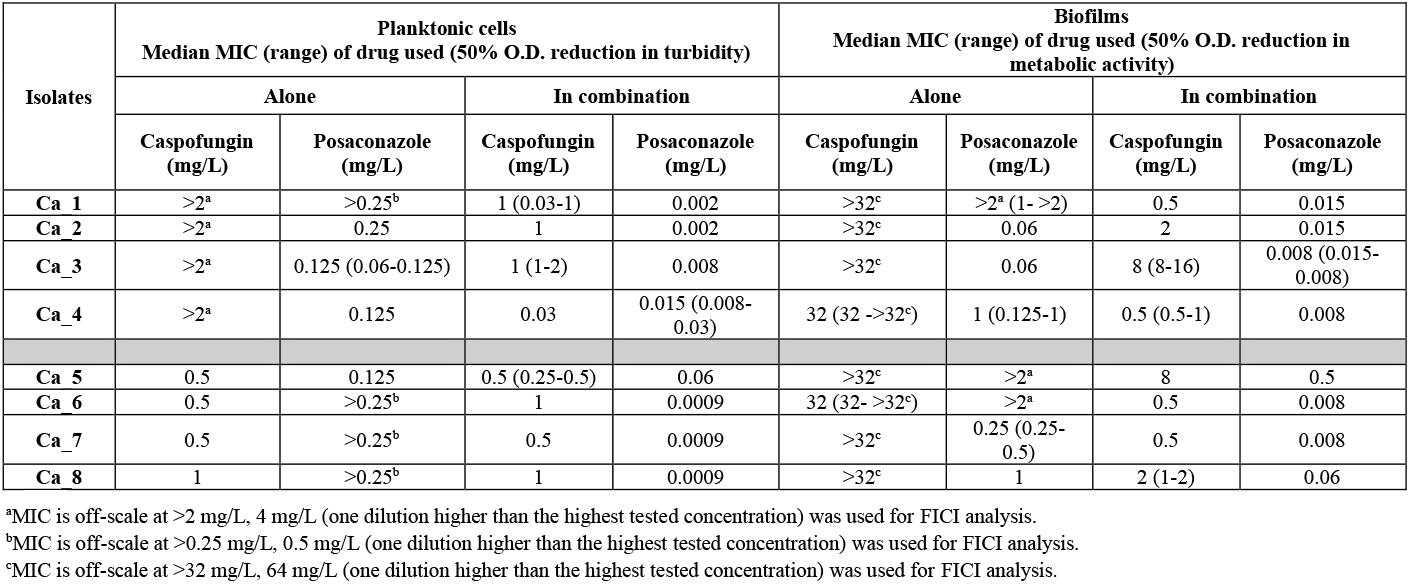
Minimum inhibitory concentrations (MICs) of caspofungin alone and in combination with posaconazole against *Candida auris* planktonic cells and biofilms.

| Isolates | Planktonic cells<br>Median MIC (range) of drug used (50% O.D. reduction in turbidity) |  |  |  | Biofilms<br>Median MIC (range) of drug used (50% O.D. reduction in metabolic activity) |  |  |  |
| --- | --- | --- | --- | --- | --- | --- | --- | --- |
|  | Alone |  | In combination |  | Alone |  | In combination |  |
|  | Caspofungin (mg/L) | Posaconazole (mg/L) | Caspofungin (mg/L) | Posaconazole (mg/L) | Caspofungin (mg/L) | Posaconazole (mg/L) | Caspofungin (mg/L) | Posaconazole (mg/L) |
| Ca_1 | >2<sup>a</sup> | >0.25<sup>b</sup> | 1 (0.03-1) | 0.002 | >32<sup>c</sup> | >2<sup>a</sup> (1- >2) | 0.5 | 0.015 |
| Ca_2 | >2<sup>a</sup> | 0.25 | 1 | 0.002 | >32<sup>c</sup> | 0.06 | 2 | 0.015 |
| Ca_3 | >2<sup>a</sup> | 0.125 (0.06-0.125) | 1 (1-2) | 0.008 | >32<sup>c</sup> | 0.06 | 8 (8-16) | 0.008 (0.015-0.008) |
| Ca_4 | >2<sup>a</sup> | 0.125 | 0.03 | 0.015 (0.008-0.03) | 32 (32- >32<sup>c</sup>) | 1 (0.125-1) | 0.5 (0.5-1) | 0.008 |
| Ca_5 | 0.5 | 0.125 | 0.5 (0.25-0.5) | 0.06 | >32<sup>c</sup> | >2<sup>a</sup> | 8 | 0.5 |
| Ca_6 | 0.5 | >0.25<sup>b</sup> | 1 | 0.0009 | 32 (32- >32<sup>c</sup>) | >2<sup>a</sup> | 0.5 | 0.008 |
| Ca_7 | 0.5 | >0.25<sup>b</sup> | 0.5 | 0.0009 | >32<sup>c</sup> | 0.25 (0.25-0.5) | 0.5 | 0.008 |
| Ca_8 | 1 | >0.25<sup>b</sup> | 1 | 0.0009 | >32<sup>c</sup> | 1 | 2 (1-2) | 0.06 |

<sup>a</sup>MIC is off-scale at >2 mg/L, 4 mg/L (one dilution higher than the highest tested concentration) was used for FICI analysis. <sup>b</sup>MIC is off-scale at >0.25 mg/L, 0.5 mg/L (one dilution higher than the highest tested concentration) was used for FICI analysis. <sup>c</sup>MIC is off-scale at >32 mg/L, 64 mg/L (one dilution higher than the highest tested concentration) was used for FICI analysis.

Table 3 summarises the *in vitro* interactions between caspofungin and posaconazole based on the median FICIs. Antagonistic interactions were never observed (all FICIs ≤4). Using a two - dimensional broth microdilution checkerboard assay and FICI calculation, the nature of the caspofungin-posaconazole interaction was found to be synergistic in the case of echinocandin-resistant isolates, both for planktonic cells and biofilms, with median FICIs from 0.247 to 0.49 and from 0.091 to 0.5, respectively. In the case of echinocandin-susceptible isolates, synergistic interactions were observed exclusively for sessile cells, with median FICIs from 0.033 to 0.375, while the nature of the interaction of their planktonic forms was indifferent, with median FICIs ranging from 1.002 to 2.001 (Table 3).

**Table 3.**
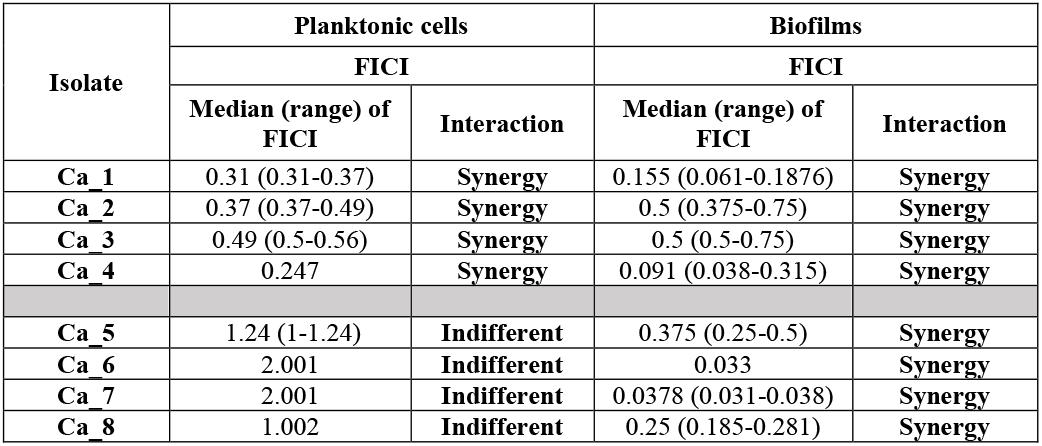
*In vitro* interactions by FIC indices (FICI) of caspofungin in combination with posaconazole against *Candida auris* planktonic cells and biofilms.

Figure 1 shows the dose-response surfaces for caspofungin–posaconazole calculations with MacSynergy II. Based on the cumulative log volumes obtained, the combination of caspofungin and posaconazole produced a moderate synergy for echinocandin-susceptible strains, with 5.71 and 6.96 cumulative synergy log volumes for planktonic and sessile cells, respectively (Figure 1A and C). In the case of *FKS* mutant isolates, a strong synergy was observed, with 16.59 and 32.39 cumulative synergy log volumes for planktonic and sessile cells, respectively (Figure 1B and D).

**Figure 1.**
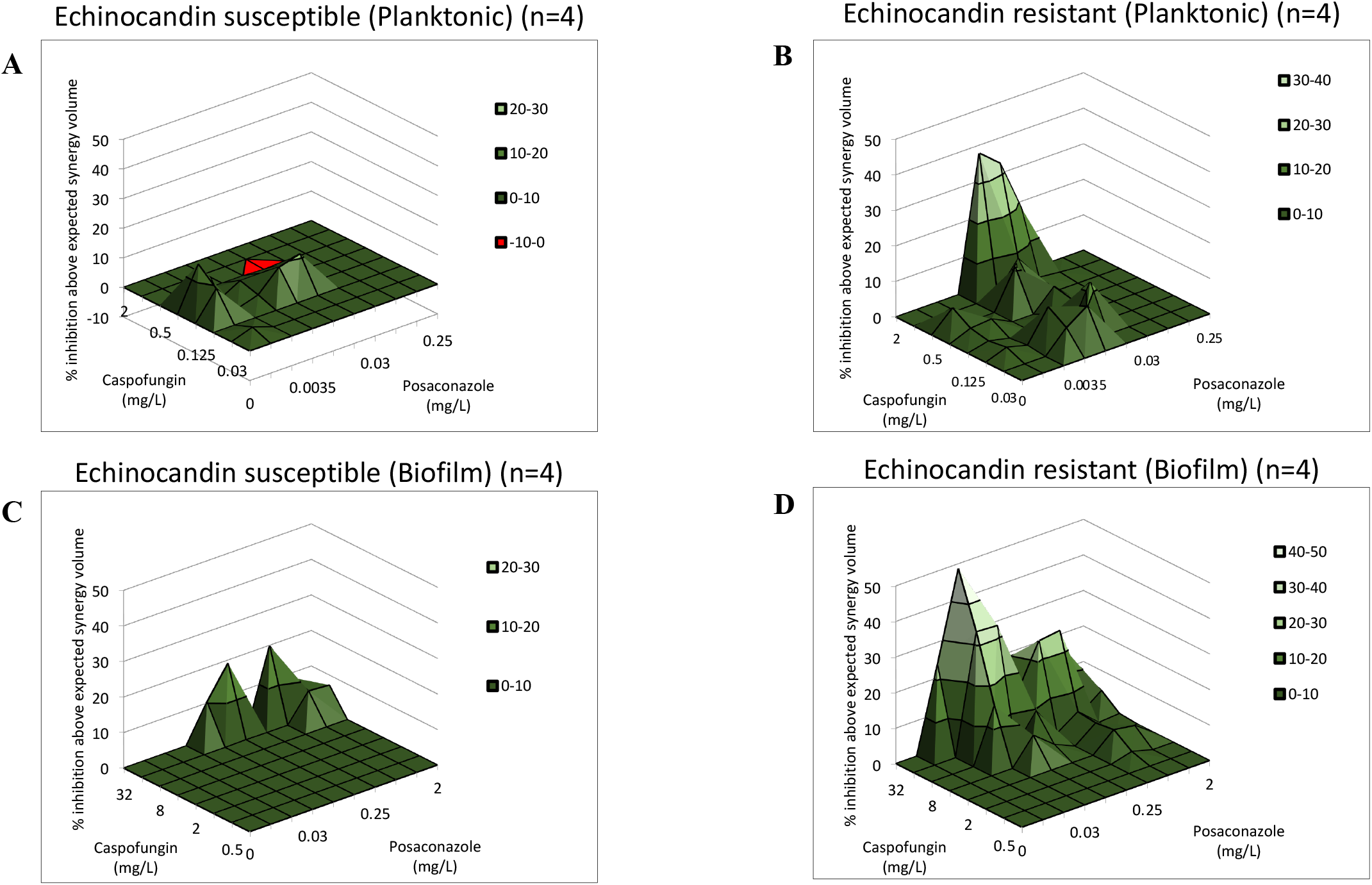
Effect of caspofungin in combination with posaconazole against *Candida auris* planktonic cells (A–B) and biofilms (C–D) using MacSynergy II analysis. Additive interactions appear as a horizontal plane at 0% inhibition. The interaction is defined as synergistic if the obtained surface is greater compared to the predicted additive surface. The volumes are calculated at the 95% confidence interval. The figures represent the cumulative synergy volume in case of four-four *FKS* wild type (A and C) and mutant (B and D) isolates for planktonic cells and biofilms, respectively.

The strong anti-biofilm effect of the combinations was confirmed by LIVE/DEAD viability staining (Figure 2). The caspofungin-exposed (4 mg/L) *C. auris* biofilms exhibited increased cell death in the presence of posaconazole (0.03 mg/L) (Figure 2D and H) compared to untreated (Figure 2A and E), caspofungin-exposed (Figure 2B and F) and posaconazole-treated sessile cells (Figure 2C and G). The disrupted biofilm structure and increase in dead cells could be observed in echinocandin-susceptible and echinocandin-resistant isolates (Figure 2D and H).

**Figure 2.**
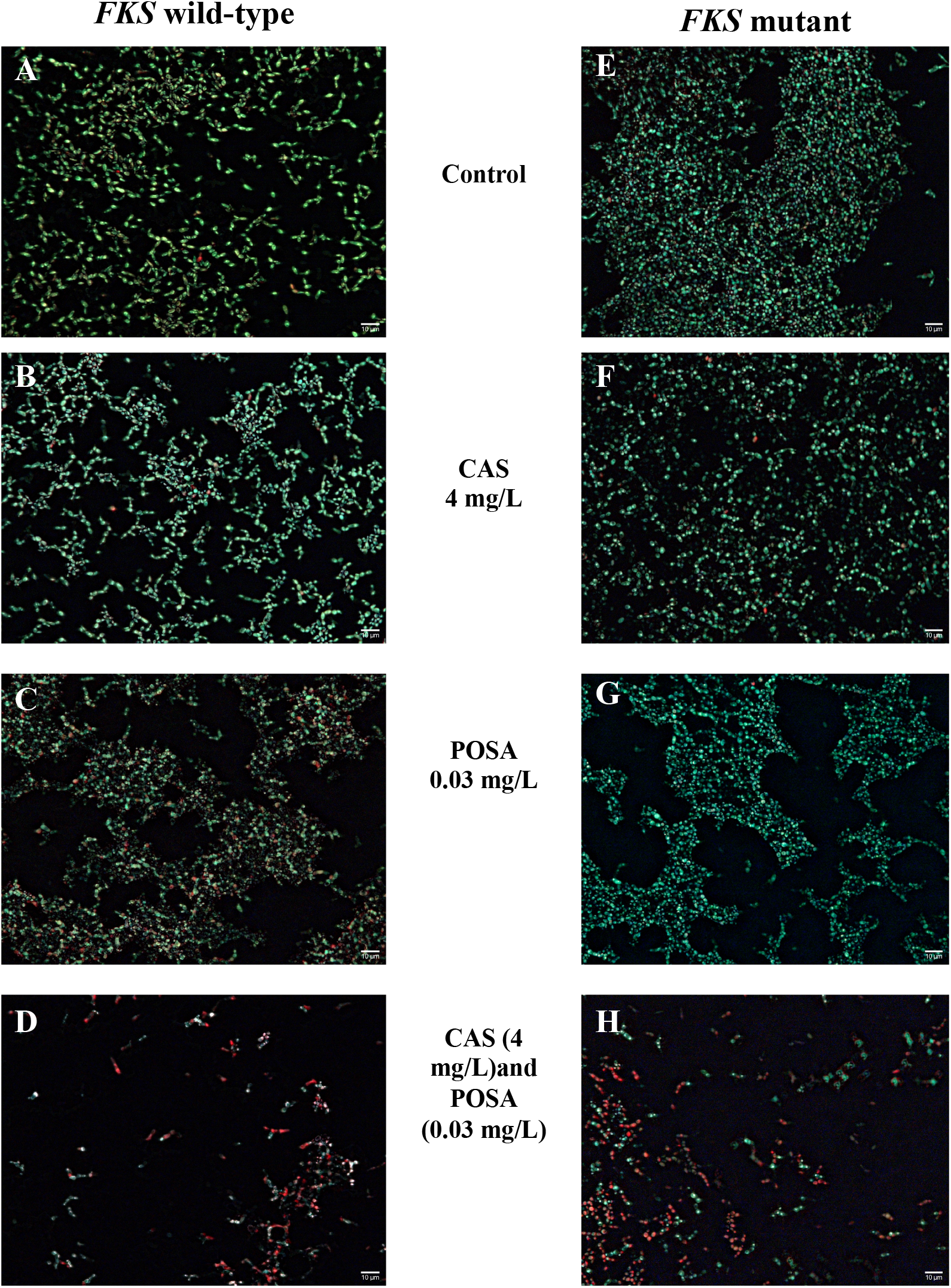
LIVE/DEAD fluorescence imaging of one *Candida auris FKS* wild-type (Ca_5) (A–D) and one *FKS* mutant (Ca_1) (E–H) representative isolates after caspofungin exposure with (D and H) and without (B and F) posaconazole. Pictures A and E show the untreated *C. auris* and biofilms respectively, while picture C and G demonstrate the posaconazole-exposed biofilms for wild-type (C) and mutant (G) respectively. Live cells (green) and nonviable cells (red) were stained with Syto9 and propidium iodide respectively. All images show typical fields of view. Scale bars represent 10 μm.

## 4. Discussion

The clinical microbiology community is increasingly reporting an alarming rise in the incidence and spread of drug resistant *C. auris*-related cases globally, which are associated with high mortality rates (30-60%) (Chowdhary *et al.* 2018, van Schalkwyk *et al.* 2019). Several current studies have provided extensive documentation of the global antifungal resistance profiles of *C. auris* isolates to azoles and echinocandins (Szekely *et al.* 2019, Umamaheshwari *et al.* 2021, Maphanga *et al.* 2021). The need for combination-based antifungal therapy against *C. auris* stems from the continuous risk of invasive infections, especially in vulnerable immunosuppressed patient populations (Vitale 2021, Bandara and Samaranayake 2022). Moreover, the biofilm forming ability of this species within these susceptible patient groups further exacerbates the risk (Sherry *et al.* 2017, Horton and Nett 2020). Despite the need for an effective and reliable approach to treatment, the clinical practice guidelines of the European Society for Clinical Microbiology and Infectious Diseases (ESCMID), the European Confederation of Medical Mycology (ECMM) and the Infection Disease Society of America (IDSA) still recommend echinocandin-based monotherapy for the majority of these infections (Arendrup *et al.* 2014, Pappas *et al.* 2016). The emergence of echinocandin resistance is therefore a relevant concern (O’Brien *et al.* 2020, Jacobs *et al.* 2022). Based on the cutoff values suggested by the CDC, the resistance rate to echinocandins is approximately 5% (Arendrup *et al.* 2017). Nevertheless, Kathuria *et al.* (2015) reported that 33 *C. auris* isolates out of 102 showed elevated MIC values (≥1 mg/L) to caspofungin in India. Furthermore, none of the echinocandins had any activity in 8% of the isolates, with MICs ranging from 4 to >8 mg/L (Kathuria *et al.* 2015). The reduced susceptibility to echinocandins is associated with mutations in the hot spot regions of the *FKS1* or *FKS2* genes (Wiederhold 2016). In a previous study, poor therapeutic response was linked to the presence of the S639F *FKS1* mutation in a systemic murine model (Kordalewska *et al.* 2018). In addition, Sharma *et al.* (2022) observed that the *FKS1* genotype was a more accurate predictor of *in vivo* response than the MIC values of the isolates. The *FKS* point mutations described in our study were detected previously (S639Y, S639P and R1354H). Al-Obaid *et al.* (2022) reported that isolates containing the S639Y or S639P mutation in hot spot 1 of *FKS1* exhibited reduced susceptibility to echinocandins, especially against micafungin. Asadzadeh *et al.* (2022) also described isolates with decreased sensitivity to echinocandins carrying a R1354H mutation in hot spot 2 of *FKS1.*

Several *in vitro* and *in vivo* studies on antifungal drugs have shown that combinations can broaden the coverage, increase the fungicidal effect in unresponsive cases and significantly decrease the risk of the emergence of acquired resistance (Vitale 2021). In addition, combination based therapeutic approaches besides monotherapy are also recommended in situations such as those depending on the type and site of infection and the patient conditions (Vitale 2021). Several studies have reported the negligible effect exerted by echinocandins in monotherapy against *C. auris,* both *in vitro* and *in vivo* (Kordalewska *et al.* 2018, Tóth *et al.* 2020, Nagy *et al.* 2021, Sharma *et al.* 2022). Regarding echinocandin-based combinations, Katragkou *et al*. (2017) showed synergistic interactions between isavuconazole and micafungin against *C. albicans, C. parapsilosis* and *C. krusei,* with the degree of synergy ranging from 1.8 to 16.7%. Fakhim *et al.* (2017) also observed synergistic interactions between micafungin and voriconazole, with FICIs of 0.15 to 0.5. In a recent study examining 36 *C. auris* clinical isolates, synergy or partial synergy was observed in 14% and 61% of the isolates, respectively, with the combination of anidulafungin and voriconazole, and in 31% and 53% of isolates, respectively, with the combination of anidulafungin and isavuconazole. Caballero *et al*. (2021) found that isavuconazole–echinocandin combinations were more effective than monotherapy regimens. These findings coincide with the results reported by Nagy *et al*. (2022), where caspofungin and isavuconazole showed a synergistic interaction in 61% of tested planktonic isolates, while the ratio was 86% in the case of one-day-old biofilms.

Previous studies have revealed posaconazole to be the most active azole, followed by isavuconazole and itraconazole, with geometric mean MICs of 0.053 mg/L, 0.066 mg/L and 0.157 mg/L, respectively (Ruiz-Gaitán *et al.* 2019). Regarding the various azoles, Tan *et al.* (2021) observed the best *in vitro* synergy effect with minocycline against 94% of tested strains, including *C. auris*. In addition, treatment with minocycline plus posaconazole significantly increased the survival of *C. auris* infected *Galleria mellonella*, where the survival rate was 51.7%. The positive effect attributed to posaconazole can also be observed in more clinical cases, where this drug was administered in combination-based therapies (Heath *et al.* 2019, Stathi *et al.* 2019, Shaukat *et al.* 2020).

An important strength of this study is that certain tested isolates have proven *FKS* mutations, which can be examined in terms of the planktonic and biofilm susceptibility to posaconazole and caspofungin in combination. Furthermore, whole genome sequencing was performed in the case of all isolates tested. Nevertheless, it should be highlighted that this study had a relevant limitation, namely the low number of isolates, and we could not cover all clades in terms of *FKS* mutation; therefore, we cannot conclude clade-specific consequences.

Despite these limitations, the therapeutic potential of caspofungin and posaconazole is unquestionable, having been confirmed against biofilms, especially in the case of *FKS* mutants at clinically achievable and safe drug concentrations. This study suggests that the administration of caspofungin with posaconazole may help to expand potential treatment strategies.

## 5. Conclusion

In conclusion, our results clearly demonstrate synergistic interactions of caspofungin in combination with posaconazole against *C. auris* especially against biofilms. However, a definitive conclusion cannot be made in terms of the effect of various *FKS* mutations on clade-specific and mutation-specific susceptibility patterns unless more studies are conducted. Finally, this study has the potential to be a starting point for further studies exploring the *in vivo* impact of these combinations against *C. auris*.

## Data availability statement

The data shown and discussed in this paper have been deposited in the NCBI GenBank with the following BioProject no.: PRJNA865124.

## Ethical approval

Not required

## Author contributions

RK conceived the ideas, analyzed the data, and wrote the manuscript. NB, FK, BB, ÁJ and AB performed the biofilm forming related tests and susceptibility tests, analysis and wrote the manuscript. ZT performed the microscope related experiments. AMB provided the strains and interpreted the obtained results, revised the manuscript. LM analyzed the data, provided sources and revised the manuscript.

